# Quantification and uncertainty of root growth stimulation by elevated CO_2_ in mature temperate deciduous forest

**DOI:** 10.1101/2021.04.15.440027

**Authors:** Clare Ziegler, Aleksandra Kulawska, Angeliki Kourmouli, Liz Hamilton, Zongbo Shi, A. Rob MacKenzie, Rosemary J. Dyson, Iain G. Johnston

**Affiliations:** Birmingham Institute of Forest Research, University of Birmingham, Birmingham, UK; School of Biosciences, University of Birmingham, Birmingham, UK; School of Geography, Earth and Environmental Sciences, University of Birmingham, Birmingham, UK; School of Mathematics, University of Birmingham, Birmingham, UK; Department of Mathematics, University of Bergen, Bergen, Norway; Computational Biology Unit, University of Bergen, Bergen, Norway

## Abstract

Increasing CO_2_ levels are a major global challenge, and the extent to which increasing anthropogenic CO_2_ emissions can be mitigated by natural carbon sinks remains poorly understood. The uptake of elevated CO_2_ (eCO_2_) by the terrestrial biosphere, and subsequent sequestration as biomass in ecosystems, may act as a negative feedback in the carbon budget, but remains hard to quantify in natural ecosystems. Here, we combine large-scale field observations of fine root stocks and flows, derived from belowground imaging and soil cores, with image analysis, stochastic modelling, and statistical inference, to elucidate belowground root dynamics in a mature temperate deciduous forest under free-air CO_2_ enrichment to 150ppm above ambient levels. Using over 67*k* frames of belowground observation, we observe that eCO_2_ leads to relatively faster root production (a peak volume fold change of 4.52 ± 0.44 eCO_2_ versus 2.58 ± 0.21 control). We identify an increase in existing root elongation relative to root mass decay as the likely causal mechanism for this acceleration. Direct physical analysis of biomass and width measurements from 552 root systems recovered from soil cores support this picture, with lengths and widths of fine roots significantly increasing under eCO_2_. We use dynamic measurements to estimate fine root contributions to net primary productivity, finding an increase under eCO_2_, with an estimated mean annual 204 ± 93 g dw m^−2^yr^−1^ eCO_2_ versus 140 ± 60 g dw m^−2^ yr^−1^ control. We also quantify and discuss the uncertainties in such productivity measurements. This multi-faceted approach thus sheds quantitative light on the challenging characterisation of the eCO_2_ response of root biomass in mature temperate forests.

## Introduction

Human-induced carbon dioxide (CO_2_) emissions are a major contributor to climate change, making up the majority of anthropogenic greenhouse gas emissions [1]. Rising atmospheric levels of CO_2_ due to anthropogenic emissions are partly mitigated by terrestrial and marine carbon sinks which take up CO_2_. However, the behaviour of land-based carbon sinks as CO_2_ levels continue to increase remains poorly understood, challenging both our fundamental scientific understanding and our ability to plan strategies to combat CO_2_ increases [2].

The world’s forests are major actors in the global carbon budget [3], and it is important to understand the effects of increased CO_2_ on the world’s forests in order to make wider climate change predictions [4, 5, 6]; this necessitates an understanding of the effect of increased carbon on plant biology [7]. Aboveground processes such as photosynthesis, although logistically challenging for mature forest systems, are amenable to in-situ measurements and remote sensing. However, belowground processes are harder to measure and less frequently studied, so constitute a substantial source of uncertainty in our knowledge of the carbon budget. The production of belowground fine roots is an important player in the global carbon budget [8], suggested to be responsible for up to 33% of global NPP [9, 10], and up to a third of C and N mineralised in temperate forest soils [11].

Carbon fertilisation, where elevated CO_2_ (eCO_2_) leads to increased biomass production, may lead to belowground root mass being an increasingly important carbon sink in a high-CO_2_ world. Increasing root mass in growing forests, and stocks and flows in mature forests pushed out of equilibrium by eCO_2_, both contribute to this sink. However, increased growth under elevated carbon dioxide leads to a greater dependence on soil nutrient availability [12, 13, 14] (particularly noted in agricultural studies due to decreased nutritional value in crops [15, 16]). Nitrogen and phosphate availability appears to be of particular importance in long-term growth response [17, 18], and may become a limiting factor in non-fertilised soils. Tight coupling between nitrogen levels and phosphate mobility in soils [19] may also lead to growth becoming phosphate limited [20]. Carbon fertilisation of fine root growth may thus be limited both in magnitude and duration, possibly limiting its long term potential as a carbon sink. However, expanding root growth may allow plants to overcome some nutrient limitation effects, as new niches can be exploited. The difficulties of non-destructive observation of below-ground systems [21] mean that the extent and timescale of carbon fertilisation of belowground root mass remains challenging to quantify, particularly in mature forests.

Previous experiments have characterised the effect of eCO_2_ on specific root systems [22, 23], finding that carbon enrichment influences (and often enhances) fine root growth and increases elemental uptake through fine roots [24, 25, 26, 27, 28, 23] although data remain sparse. There is a general consensus that eCO_2_ leads to an initial increase in carbon capture and growth in both above-ground [29] [30] and below-ground tissues [31] (including in our experimental site described below [32]). Growth increases may be limited by nutrient availability [33, 34, 35], with many short-term experiments failing to capture this effect [36] which may only be visible in experiments run over many years [12]. Contributions to the experimental and theoretical challenges in the field include the twin logistic difficulties of elevating CO_2_ in a natural ecosystem and performing repeated non-invasive belowground measurements therein, the heterogeneity of natural belowground root systems, and the natural variability in root dynamics.

Here, we address these difficulties with an interdisciplinary workflow, coupling free air CO_2_ enrichment (FACE) experiments [37, 38, 39, 40] in a natural ecosystem, large-scale belowground imaging with minirhizotrons in parallel with destructive soil coring, a semi-automated image analysis pipeline, and novel applications of mathematical tools from stochastic processes and statistical inference. We work in a native, mature, deciduous UK woodland [37], motivated by the observation that climate model uncertainty in carbon-cycle feedbacks at temperate land ecosystems at these latitudes is, with equatorial land, the highest worldwide [41]. We characterise belowground root growth in this ecosystem, and show that the carbon fertilisation effect from eCO_2_ on root production is detectable, although some associated physiological changes are low in magnitude. We estimate corresponding changes in fine root contributions to net primary productivity (NPP), while quantifying and highlighting the large uncertainty that is necessarily associated with these measurements in real ecosystems. We conclude by discussing the implications of our results and analysis for natural carbon budgets under eCO_2_ and in future research in the field.

## Results

Our field observations were carried out at the Birmingham Institute of Forest Research (BIFoR) FACE facility [37] in the UK (Fig. 1A). The facility has been built into a native mature deciduous woodland, dominated by oak (*Quercus robur*) interspersed with overstood hazel (*Corylus avellana*) coppice. Sycamore (*Acer pseudplatanus*), hawthorn (*Crataegus monogyna*) and holly (*Ilex aquifolium*) have self-seeded into gaps and, with the hazel, form a distinct sub-canopy. The forest grows on centuries-old Orthic Luvisol soil with a mulmoder humus classification [37, 42]. The experimental design consists of three eCO_2_ and three ambient-air control regions (as in the Australian EucFACE study [43]). These regions, which we refer to as arrays, are 30m diameter rings, with free-standing towers extending above the”25m oak canopy (Fig. 1B). Pipes attached to the towers emit treated air with increased CO_2_, in order to raise the CO_2_ levels within the eCO_2_ arrays to 150 ppm above the ambient (calculated as the lowest CO_2_ mixing ratio measured in the control arrays). The control arrays are identical, but the air released into the array contains no additional CO_2_. Performance of the facility was excellent over the course of these experiments, with details and further information available in Refs. [37, 32, 44].

**Figure 1:**
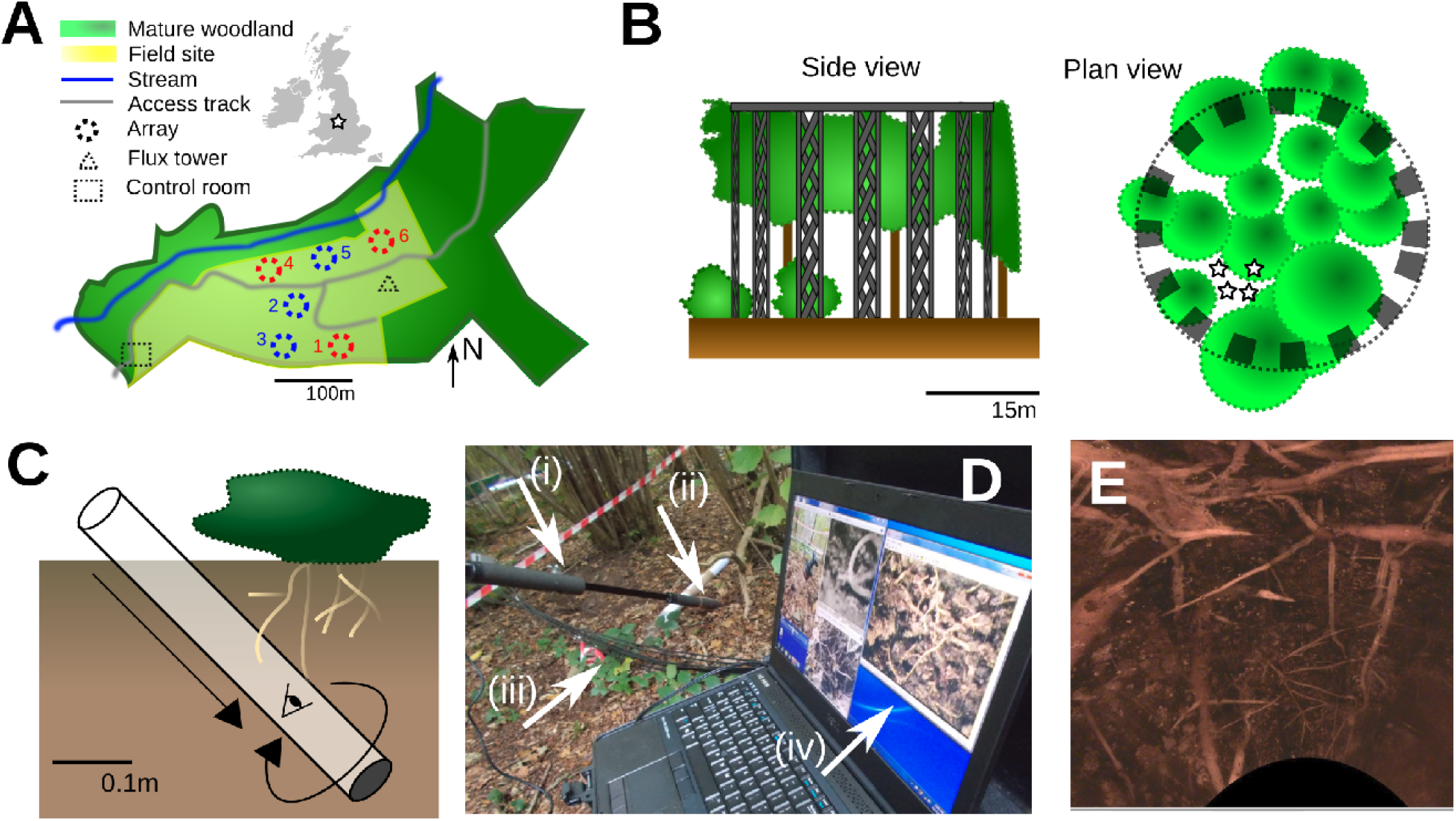
Field experiments tracking root dynamics under control and eCO_2_ conditions. **(A)** The BIFoR FACE experimental facility is situated in mature deciduous woodland near Stafford, UK, and contains three eCO_2_ (red) and three control (blue) ‘arrays’. **(B)** Arrays consist of scaffolding (grey) supporting pipes carrying eCO_2_ or ambient air to the forest canopy. Each array has four minirhizotron installation sites. (A-B) adapted from Ref. [37]. **(C)** Minirhizotron sites consist of a transparent tube embedded at a 45° angle in the soil, covered and sealed when not in use. **(D)** A camera and lighting system (i) is inserted into each of these tubes (ii) to take belowground images of *in situ* root systems around the tube circumference and along its length. The imaging system is connected (iii) to a field power supply and to a computer running real-time image acquisition software (iv). **(E)** Illustration of data acquired from minirhizotrons; this composite consists of many images concatenated and embedded on a cylindrical manifold for illustrative purposes.

BIFoR FACE, situated in a mixed woodland, involves studying plants within a diverse preexisting ecosystem rather than a plantation as with many previous FACE experiments [45, 46]. Therefore, data collection methods need to be as non-destructive as possible. Minirhizotrons allow a small subset of a root system to be observed over time [47], but can by nature obstruct the natural structure of the root system, and require indirect biomass quantification [48]. Soil cores are a more destructive sampling method and do not allow long-timescale observation of the same region, but allow for more direct estimations of root biomass and turnover [49]. Both methods have been successfully employed in previous studies of fine root growth [25]. Here, we use a workflow coupling the two modes of investigation (see Discussion) with new theoretical developments and a rigorous treatment of uncertainty [50] to maximise the interpretability and transferrability of our results.

### Fine root structure observed with minirhizotrons

We used minirhizotrons (Fig. 1C; see Methods) to observe belowground root systems (Fig. 1C-E; Fig. S1) in control and eCO_2_ plots across a two-year time frame. 216 eCO_2_ and 216 control measurements were taken, each involving around 150 individual frames of belowground imaging, for a total 6.7 × 10^4^ frames of observed root volumes. To facilitate working with this data volume, we designed and used a semi-automated image analysis pipeline (see Methods) to quantify observed root dry weight biomass. The root volume data showed a pronounced seasonal trend (Fig. 2A), with both eCO_2_ and control volume increasing from April to October 2017, then decreasing until June 2018 before increasing again through the late summer. In addition to general seasonal variation, these trends likely also include some system-specific influences. First, some of the earliest seasonal change may be due to a wounding response, inducing increased root production, due to the disturbance caused by the minirhizotron installation four months earlier (see Methods). Second, some of the reduction in biomass across the arrays in the second year of sampling may be attributed to environmental effects on the forest. The summer of 2018 was particularly dry, and the trees were recovering from a springtime infestation of winter moth (*Operophtera brumata*) caterpillars (17th May to 3rd June) [51]. These influences demonstrate the importance of the control measurement set, characterising the seasonal and specific variation experienced by the system and allowing us to account for additional behaviour due to eCO_2_.

**Figure 2:**
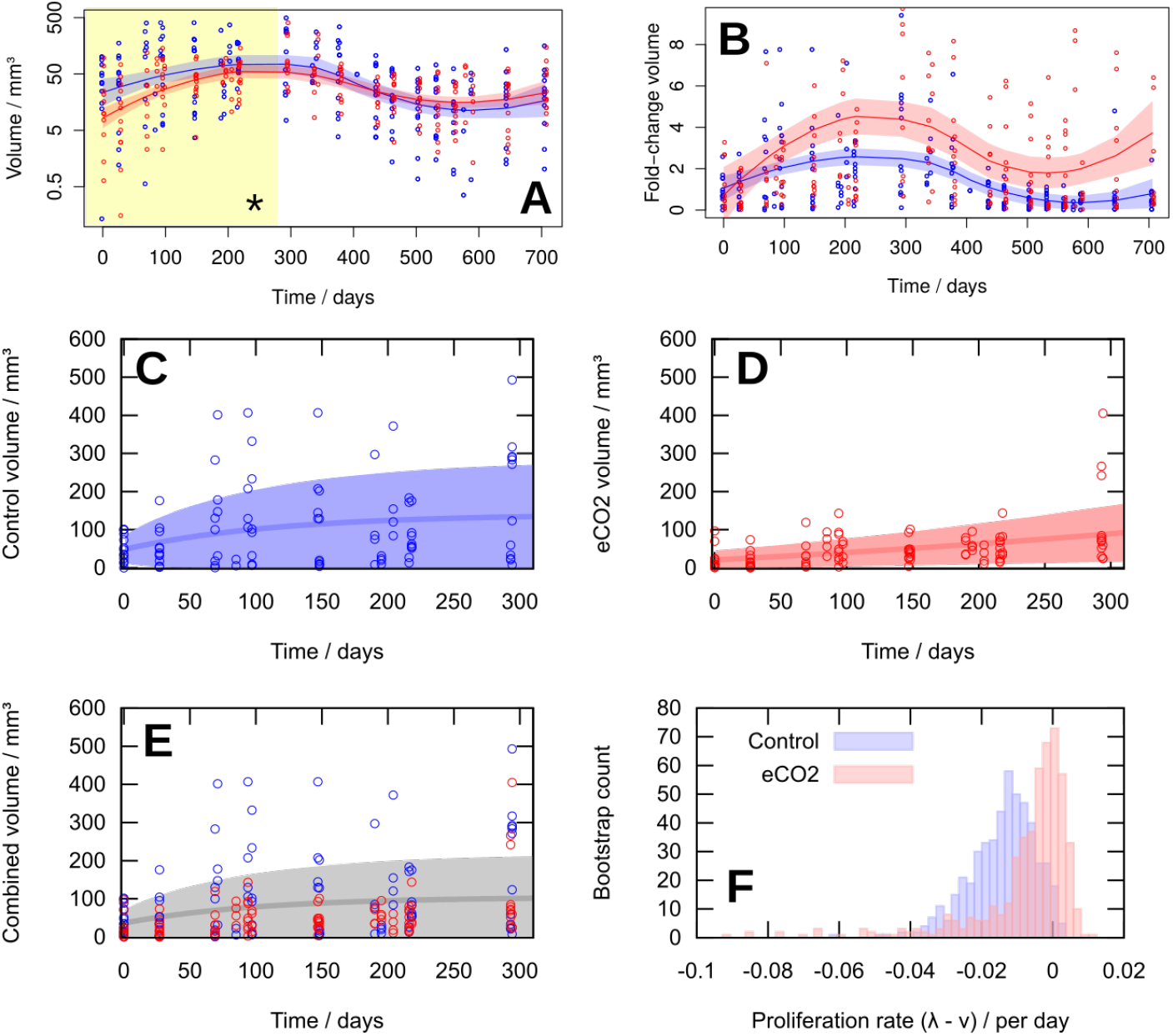
Root volume changes over time under eCO_2_ and control conditions. **(A)** Fine root volume observations in eCO_2_ (red) and control (blue) experiments. Each datapoint is an observation from a single minirhizotron site. LOESS fits to the log-transformed data are shown with 95% confidence intervals, displaying the different rates in the first 300-day period (*, highlighted). **(B)** Fold change in fine root volume from initial observations. Each datapoint is an observation from a single minirhizotron site, normalised by the initial volume averaged across all sites in an individual array. **(C-E)** The birth-immigration-death (BID) model described in the text, applied to the first year of (C) eCO_2_, (D) control, and (E) combined volume observations. Time axis gives days from 11 April 2017. The maximum likelihood BID parameterisation is found for each dataset, then the mean and standard deviation of the model for that parameterisation is plotted. A likelihood ratio test shows statistical support for the individual models (C)+(D) over the combined model (E) (*p* < 10^−15^). **(F)** Bootstrapped estimates for the difference between root elongation (birth, λ) and root decay (death, *ν*) parameters for eCO_2_ and control data. λ – *ν* is higher (with positive maximum likelihood estimate) for eCO_2_, reflecting increasing root proliferation, and lower (with negative maximum likelihood estimate) for control, reflecting decreasing proliferation. Data points are measurements from individual minirhiztoron tubes.

The average absolute magnitudes of root volumes under eCO_2_ and control were broadly similar, but the rates of increase of volume in spring-summer 2017 (and late summer 2018) appeared higher for eCO_2_ than for control. To investigate this further, we calculated the fold-change increase in volume from the initial measurement for eCO_2_ and control observations (Fig. 2B) and observed dramatically higher rates of increase for eCO_2_ observations. LOESS fitting showed a maximum mean fold-change increase of 2.58 ± 0.21 for control plots and 4.53 ± 0.44 (standard errors) for eCO_2_ plots. LOESS confidence intervals cannot be directly subjected to hypothesis testing, but individual observations statistically support this difference: for example, comparing fold-change observations from 150 to 300 days into the experiment give 4.59 ± 0.58 for eCO_2_ and 2.50 ± 0.31 for control (standard errors), with *p* = 0.016 from the Mann-Whitney test.

Although suggestive of a treatment effect, these fold-change measurements must be interpreted with caution as they assign more statistical weight to the initial volume measurements, which provide a reference for the subsequent scaling. To explore these trends further, we employed a stochastic model for the belowground processes influencing fine root volume.

### Stochastic modelling for fine root volume

To make further progress, we sought a modelling framework that allowed us to harness the full distributional detail of our observational data to identify the mechanisms modulating fine root volume in control and eCO_2_ experiments.

The theory of stochastic processes provides such a framework, describing the influence of stochastic mechanisms on the probability distributions of some observed system.

Here, the observed system is the fine root volume in our experiments, and the mechanisms we consider are growth of existing roots, new roots entering the field of observation (the viewing region of a minirhizotron) and disappearance of existing roots. These mechanisms have well-studied analogues in stochastic processes: so-called birth (the replication of existing elements), immigration (the arrival of new elements from outside the system) and death (the removal of existing elements). We therefore work in a birth-immigration-death (BID) modelling framework (see Methods). The task is, given observations of fine root volume, to infer the rates of birth, immigration, and death in eCO_2_ and control experiments.

This approach has several strengths in the analysis of data like ours. First, the distributional detail of observations is captured, so that scientific information can be gained from the variance of observations as well as their mean trends. Second, the approach is naturally dynamic, allowing time series data to be naturally analysed, with more time points providing more statistical power. Third, the BID model supports both stable and exponentially-varying solutions, allowing transient as well as long-term effects to be learned from the data. This third point, in separately accounting for initial and ongoing behaviour, mitigates against the high weighting of initial observations in the fold-change calculation above.

We proceed by obtaining the likelihood function associated with the BID model (see Methods), which describes the probability of seeing a given amount of root volume at a given time when the rate parameters take given values. The parameter values that maximise this likelihood function for our data then correspond to the most likely rates for the three processes. Noting that Fig. 2A shows the strongest dynamic differences between eCO_2_ and control experiments in the first 300 days, we first analysed our observations from this time window using the BID model (see Methods). We found that the parameters inferred to describe fine root dynamics differed significantly between eCO_2_ and control experiments in the first year (likelihood ratio test, p < 10^−15^). The maximum likelihood parameterisation for control experiments supported dynamics that reached a volume steady-state, while the maximum likelihood parameterisation for eCO_2_ experiments supported an exponential increase over this period (Fig. 2C-D). The likelihood ratio test supporting the distinction between eCO_2_ and control experiments is relative to an amalgamated model with less support (Fig. 2E). This picture is supported by the maximum likelihood value of λ – *ν*, the difference between birth and death rates, which is positive (0.0045 day^−1^) for eCO_2_ experiments and negative (−0.0086 day^−1^) for control experiments. Bootstrapping with the percentile method confirmed this difference (*p* = 0.041, Fig. 2F).

The BID model applied to the second year showed no statistical support for a model where eCO_2_ and control root dynamics differed (likelihood ratio test), with limited differences in inferred parameters (Supplementary Figure S2). As described above, root growth in other time periods was lower, potentially due to environmental factors. This absence of significant differences of course cannot be interpreted as a significant absence of a difference, and perhaps the more likely explanation is that absolute growth in this period was too low for differences to be detected (see below and Discussion).

### Sampled root biomass and morphology from soil cores

In addition to minirhizotron measurements, we obtained periodic soil cores from the experimental plots and assessed the live and dead biomass in O (organic), A (mineral), and B (subsoil) horizons in these cores (see Methods). The specific depths of these horizons differed across the field site, and we found little evidence for systematic differences between horizons in live or dead biomass from soil cores (Fig. 3A-B). More biomass was often found in the lower, B horizon, reflecting the larger root structures that were sometimes sampled at this depth. The presence of less frequent but larger roots in deeper soil is also reflected in the increased variability in biomass observed in the B horizon compared to the other depths. Broadly, live biomass measurements were typically higher than dead biomass measurements.

**Figure 3:**
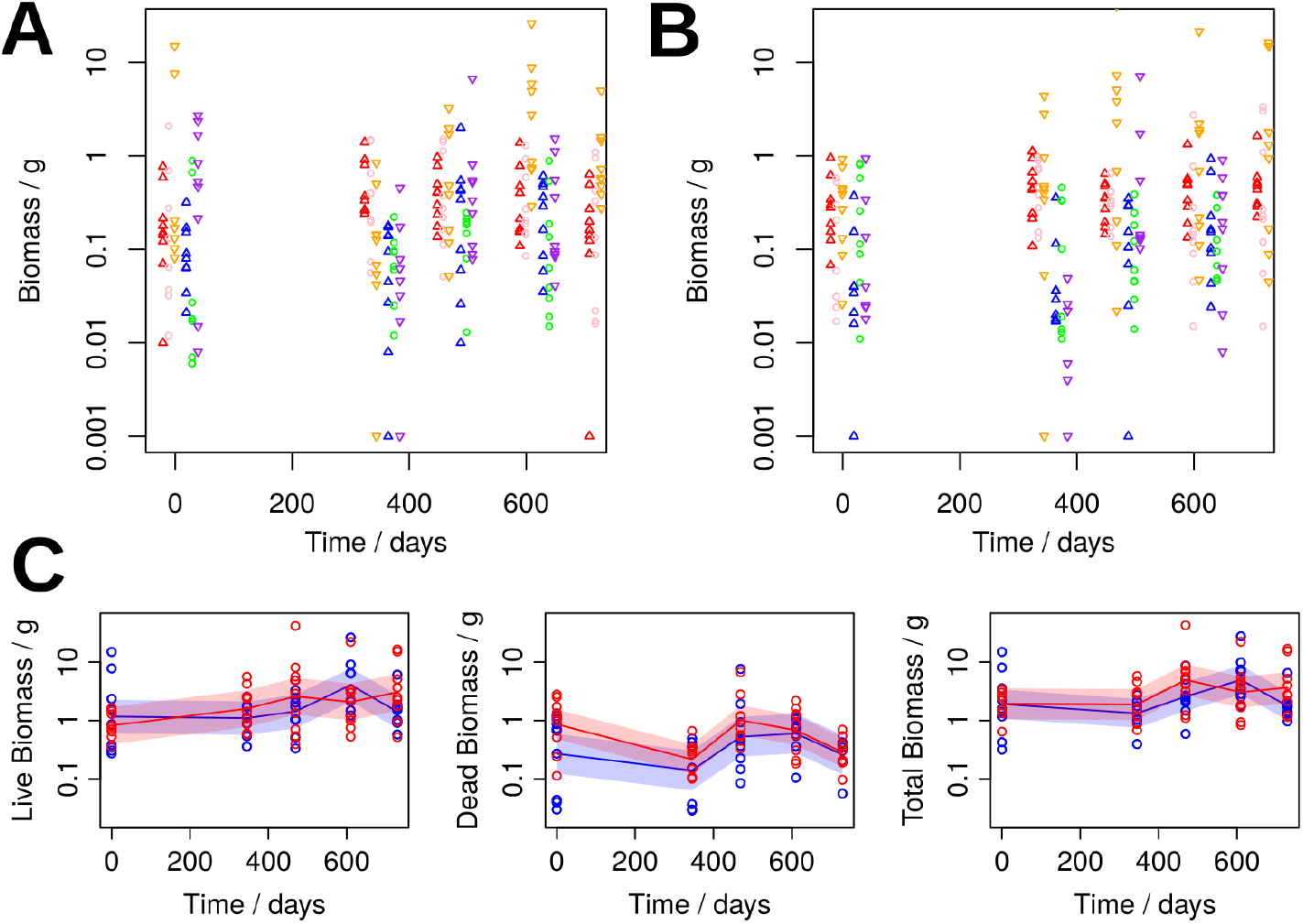
Root biomass sampled from soil cores. **(A-B)** Biomass measurements separated by soil horizon for (A) control and (B) eCO_2_ experiments. Time axis gives days from 11/04/2017. Horizons are O (up arrows), A (circles), B (down arrows); biomass is living (warm colours) and dead (cool colours). Data points are measurements from individual soil cores. **(C)** Biomass summed over all soil horizons for eCO_2_ (red) and control (blue) experiments, classified by living, dead, and total. Each datapoint corresponds to a single array. LOESS fits with 95% confidence intervals are shown.

Collecting samples across horizons (Fig. 3C), the total live biomass did not vary dramatically over the time course, although there was a trend for more rapid increase in eCO_2_ than control experiments, matching the observations from minirhizotrons.

The total dead biomass from the collected samples across horizons, however, did display some time variability, with similar trends in the eCO_2_ and control experiments (Fig. 3C). A decrease in dead biomass from April 2017 to April 2018 was observed, agreeing with the increasing observations of fine root biomass from minirhizotrons over this period. Subsequently, in the spring-summer transition in 2018, an increase in dead biomass was observed, mirroring the decrease in fine root volume observed in this period in the minirhizotrons. During and after late summer 2018, as the fine root biomass observations increase, the amount of dead biomass begins to decrease again.

Widths of roots from soil cores also displayed time variability, with root segment widths under eCO_2_ becoming greater than those in the control arrays after a year under eCO_2_ (Fig. 4A). To explore these trends further, we manually recorded the length and diameter of fine roots (< 2mm, see Methods) from 552 root systems (3709 roots in total) recovered from soil cores taken in March 2019 (Fig. 4B-D). We found a low magnitude, but statistically robust, increase in root width in eCO_2_ versus control (mean eCO_2_ 2.89 × 10^−2^ cm, mean control 2.79 × 10^−2^ cm, 1.04-fold increase), Fig. 4C). A similar signal was found in estimates of lateral root length (Supplementary Fig. S3), although these length estimates should be interpreted with some caution (see Supplementary Information). Taken together, these results suggest a detectable but slight increase in elongation and thickening of fine roots after two years under eCO_2_.

**Figure 4:**
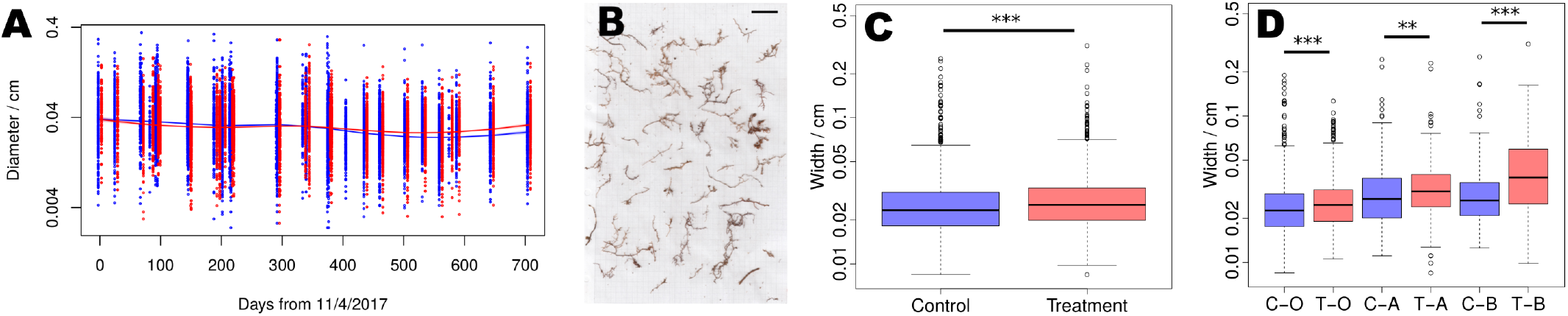
Root widths sampled from soil cores. **(A)** Fine root diameter minirhizotron observations in eCO_2_ (red) and control (blue) experiments. Each datapoint is an observation from a single root segment. LOESS fits to the log-transformed data are shown with 95% confidence intervals. **(B)** Example root system fragments from soil coring for individual analysis. Roots are shown on 1cm graph paper; scale bar shows 5cm. (C-D) Diameter of live roots taken from March 2019 soil cores under eCO_2_ (red) and control (blue) conditions. **(C)** Logged diameter of live roots separated by treatment and control. A Mann-Whitney test shows a small but significant increase in root width under eCO_2_ (1.04-fold, p = 5 × 10^−9^). **(D)** Logged root diameters from (C), separated by soil horizon for eCO_2_ (red) and control (blue). The increase in width under eCO_2_ is consistent across all 3 horizons (1.03-fold, *p* =1.5 × 10^−7^; 1.02-fold, *p* = 0.0031; 1.46-fold, *p* = 8.4 × 10^−7^ for O, A, and B respectively).

### Net primary productivity estimation and uncertainty

We next sought to estimate the contribution of root production to net primary productivity from belowground root dynamics. To this end, we follow an established method involving time-separated measurements of the same element of a root system along the top strip of the rhizotron viewing window [24, 52, 53]. Here, individual roots are identified over several monthly observations, and their dynamics are tracked over this time window (Fig. 5A). Experimental uncertainty is associated not only with observations of root structure in this protocol, but also with the estimation of mass from this structure, and the physical parameters involved in the subsequent analysis (for example, the depth of viewing window away from the minirhizotron surface).

**Figure 5:**
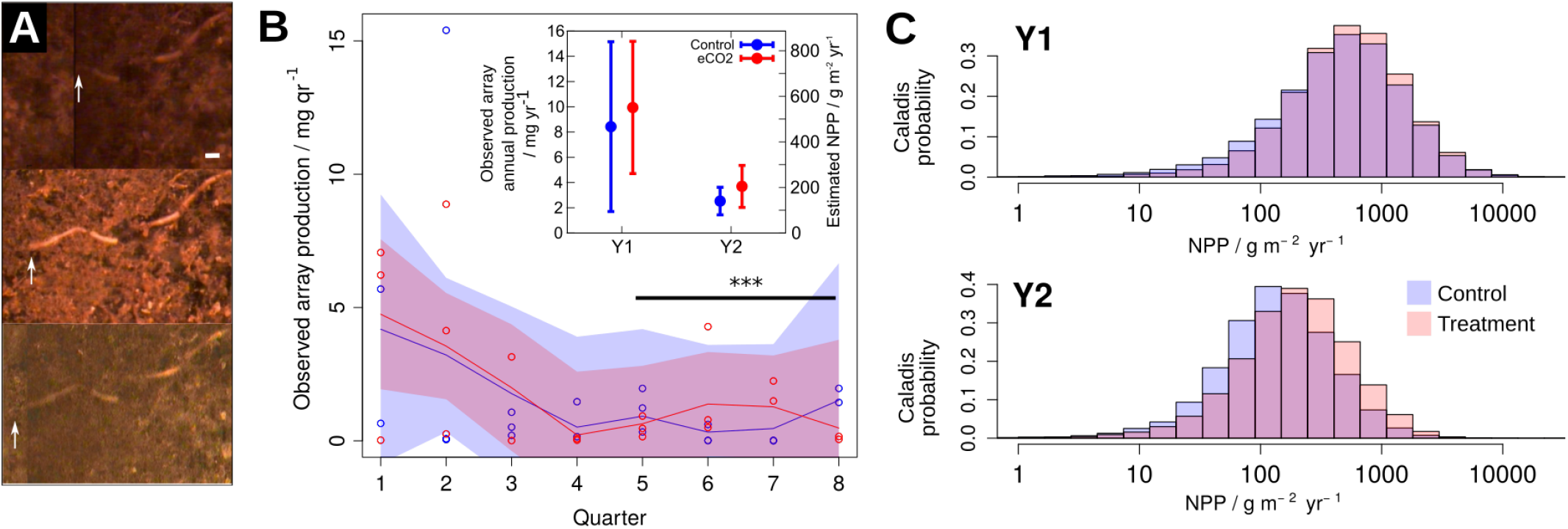
Net primary productivity estimates and uncertainties. **(A)** Example of month-by-month root dynamics (here, elongation) for NPP calculations. Arrows identify the root tip and the horizontal scale bar represents 1mm. **(B)** Observed fine root production per plot per quarter and total across plots per year (inset) from root observations under eCO_2_ (red) and ambient air (blue), with LOESS fits and 95% confidence intervals. Dynamics in quarters 5-8 (* * *) display significant differences between eCO_2_ and control (likelihood ratio test comparing quadratic models, *p* = 2.6 × 10^−4^), though see text for interpretation. **(C)** Estimates from Caladis [50] of NPP in year 1 and year 2 of our observations. Distributions reflect the uncertainty in these estimates, derived by propagating uncertainty in each value involved in the calculation.

We first characterised the increase in observed root production (ORP, the estimated increase of root biomass seen over time via minirhizotrons) in our system, before attempting to map this quantity to a readout of NPP. To calculate ORP, we multiplied observed increases in root volume *p_obs_* by estimated root density *ρ* (mass per unit volume), to estimate a biomass. Joint mass and volume measurements of sampled roots from soil cores (see Methods) yielded a mean density estimate of *ρ* = 0.34 ± 0.16 g dw cm^−3^.

We found that ORP varied substantially between arrays (Fig. 5B) but that some trends were detectable over time. First, early ORP measurements – the first six months after the experiment started – were rather higher than later measurements in both control and eCO_2_ cases. This is likely an out-of-equilibrium effect due to fine roots growing back into the field of observation of the minirhizotrons. Secondly, after these transient high values decreased, the ORP dynamics in control and eCO_2_ cases differed in quarters 5-8, with eCO_2_ ORP starting lower, increasing to a higher peak, and decreasing to a lower trough than control. This observation was supported statistically by fitting a quadratic model to the control and treatment data in both years. No significant differences were found in quarters 1-4, but in quarters 5-8 a likelihood ratio test shows significant (*p* = 2.6 × 10^−4^) support for a model where control and treatment dynamics differ (with different curvatures) over a combined model where the dynamics are the same.

Though suggestive of a higher peak fine root production in eCO_2_ conditions, this result must be interpreted with caution due to the variability and uncertainty (see below) inherent in these observations. Total annual ORP was higher in both years in eCO_2_ than control, but this coarse-grained measurement neglects the substantial within-year variability between the experiments, limiting the statistical significance of this observation.

To connect this ORP behaviour with an NPP estimate, we calculated how minirhizotron observations can be scaled to area-wide productivity estimates, while tracking uncertainty in the numerous quantities involved in this calculation. Based on the geometry of our observational setup, we derived the following expression for NPP (see Methods and Supplementary Information):

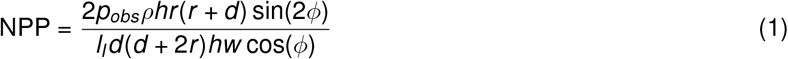

where *p_obs_ρ*, observed volume increase multiplied by estimate root density, is the ORP observed through minirhizotron samples (see Methods), *h* is the length and *w* the width of the viewing area, *r* is the radius of the minirhizotron tube, *d* is depth of viewing field, *ϕ* is the angle of the minirhizotron tube and *l_I_* is the viewing arc length. Briefly (full explanation in Methods), Eqn. 1 maps the productivity observed in the part-cylindrical viewing region of the minirhizotron tube to the corresponding surrounding volume of the soil column, and maps this volume to the corresponding 2D surface area for interpretation as a traditional NPP measurement. Without considering uncertainty (see below), the geometric aspects of Eqn. 1 provide an approximate multiplicative scaling of ~ 5.54 × 10^4^ for mapping ORP in g dw yr^−1^ to NPP in g dw m^−2^ yr^−1^.

Each parameter in Eqn. 1 has substantial associated uncertainty. To improve the interpretability of our NPP estimate, we used Caladis [50], a tool for uncertainty tracking and quantification, to propagate these uncertainties through the calculation and hence characterise the uncertainty in the final output. Expanding upon traditional uncertainty propagation, Caladis estimates full distributions over a quantity of interest given distributions over uncertain contributory factors.

Following the trends in Figs. 2–4, estimated NPP was higher under eCO_2_ in both years, albeit with large uncertainty when annual summaries are used. Specifically, estimated mean NPP values in year 1 were 467 ± 372 g dw m^−2^ yr^−1^ for control and 551 ± 290 g dw m^−2^ yr^−1^ for eCO_2_ (1.17-fold higher mean with eCO_2_), and in year 2 – where transient behaviour plays a less pronounced role – were 140±60 g dw m^−2^ yr^−1^ for control and 204±93 g dw m^−2^ yr^−1^ for eCO_2_ (1.45-fold higher mean with eCO_2_). The mapping from ORP to NPP substantially increases the uncertainty on these results: the distributions associated with the overall estimates are shown in Fig. 5C. This approach to calculating NPP thus characterises the very substantial uncertainty involved in estimating NPP from minirhizotron measurements. We suggest that care should be taken when interpreting NPP results, which are likely to involve ‘hidden’ uncertainties from the way the estimates are constructed.

The period where productivity differences between control and eCO_2_ experiments are most pronounced coincides with the previous observed increases in root width and length (Fig. 4), and not with the earlier increase in root proliferation (Fig. 2). Taken together, these observations suggest a picture where eCO_2_ supports faster proliferation as roots are expanding into unoccupied space, and increased productivity due to larger fine roots in a more stable state.

## Discussion

We have used a novel combination of field experiments and theoretical approaches to explore the dynamics of belowground fine root growth in eCO_2_ conditions, in a natural mature temperate deciduous woodland – a class of ecosystem that is both of central importance in global carbon budgets [41] and which reflects a substantial source of uncertainty in current understanding of carbon feedbacks [41]. Our multifaceted, interdisciplinary approach both allows a dissection of the mechanisms involved in eCO_2_ responses and a rigorous quantification of uncertainties, rates, and timescales of these processes.

Taken together, our results suggest that eCO_2_ provides a detectable stimulation of belowground root growth in our mature temperate woodland. We found that relative rates of fine root proliferation detectably increased under eCO_2_ in the first year of treatment and also differed, to a lesser extent, in the second year. Subsequently, the summer of 2018 (July to September) was particularly dry [44] and the trees were recovering from a springtime winter moth (*Operophtera brumata*) caterpillar infestation (17th May to 3rd June) [51], both of which features may have challenged root proliferation. Some increased growth in the first year could also be attributed to a wounding response due to disturbance from minirhizotron installation four months earlier. This response is clearly amplified due to eCO_2_, but may not reflect steady state behaviour. At later stages of the experiment, other differences became clear, with increases in fine root width and length in later soil cores revealed through large-scale morphological assessment, and an increase in observed fine root production under eCO_2_. This pattern of observations is compatible with a picture where eCO_2_ has joint effects, which the different years of our observations illustrate: during expansion, when roots are growing into new space, eCO_2_ increases the proliferation rate of fine roots, and in more stable root layouts, eCO_2_ supports the expansion of existing roots.

The magnitudes of the effects we observe under eCO_2_ vary by observation. Some of the increases in root morphological characteristics we observe are rather small-scale (the small associated p-values are due to the large volume of observational data we obtained rather than the magnitude of the effect). Changes in root elongation, and in observed root production and estimated NPP, particularly in the second year, are of rather higher magnitude, though subject to more uncertainty. The largest effect we observe is the increase in peak observed root productivity in the second year, suggesting that the increase in root size due to eCO_2_ fertilisation may be the dominant effect in our system. Considering means alone, eCO_2_ was responsible for a 46% increase in annual estimated NPP, reflecting a potentially substantial influence on the belowground system – as found in studies of other systems [24, 25, 26, 27]. However, the dynamics leading to these results are nuanced and the values are subject to uncertainty, as described above, and hence propagating such values into other models and analyses must be done with caution and this uncertainty explicitly tracked.

As with any experimental approaches aimed at this challenging system, the methods we use require some scrutiny. For comparison, we first note that our estimates of NPP are broadly similar to those from similar studies (which fall, for example, around 100-200 g dw m^−2^ yr^−1^ [27]; 100-400 g C m^−2^ yr^−1^ [24], and 40-500 g C m^−2^ yr^−1^ in Wytham woods, a more similar ecosystem [54]). Observation of year-to-year variation in this behaviour is not inconsistent with other studies [26]. We note here that the conversion from estimates of dry weight biomass (g dw m^−2^ yr^−1^) to carbon biomass (g C m^−2^ yr^−1^) includes another uncertainty, the proportional contribution of carbon to root dry mass. We do not focus on this conversion here but a convenient heuristic is that dry mass is around 50% carbon [55].

Our approach combines methods to avoid particular reliance on one experimental technique. Minirhizotrons have been used for many years to measure fine root productivity and turnover [56], and are superior for the monitoring of very fine roots than sequential soil coring, which can lead to the loss of fine root matter during processing [57, 58], although more data is available from soil core studies [59]. Soil cores are useful for the quantification of root standing stock [60, 61, 53], and are particularly powerful when combined with minirhizotrons [62, 63]. Ingrowth cores are often used in similar studies, but are unsuitable for sites with strong seasonal fluctuations [64, 65]. However, minirhizotrons present their own challenges to the researcher [22, 66], and careful consideration is required to produce accurate results [67]. Careful installation is required to minimise soil compaction and air gaps [48], and variations in depth of field between soil types may lead to underestimation of root width [68]. However, carefully collected and interpreted fine root data is essential to our understanding of carbon cycling and forestry processes [69, 70].

One source of variability in our study is the diversity of plant life in our research site. We do not attempt to classify roots based on phylogenetics, instead relying on our sampling across physical positions and different arrays to mitigate against any systematic bias towards particular species. This is supported by the consistent trends we observe through several different modes of observation, and means that our results should be viewed as ecosystem-wide readouts rather than species-specific responses.

To conclude, we believe that our multi-faceted approach, using large volumes of diverse data and tracking uncertainty throughout scaling calculations, is a powerful way to address challenging questions about hard-to-observe ecosystem responses. The use of stochastic modelling rather than purely data-driven analysis increases our approach’s power to detect mechanistic differences, and we believe our consideration of the (large) uncertainties involved in belowground observation has helped increase the interpretability of our findings by underlining the necessary caution required [50]. We hope that these approaches help inform the quantitative interpretation of past and future experiments on this globally important topic.

## Methods

### Minirhizotron installation

Four 50cm long minirhizotron tubes sealed with bungs (custom machined at University of Birmingham) were installed in each array, with the sites chosen to keep the species makeup of surrounding vegetation as consistent as possible. A Van Walt (Prestwick Lane, Surrey, GU27 2DU) 55mm corer was used to remove a cylinder of soil at a 45 degree angle. The minirhizotron tube, sealed at the lower end and with a removable bung at the upper end, was then inserted manually into the hole, with care taken to prevent damage to the exterior of the tube. Each set of four was installed in a designated area within the array with clear markers to minimise footfall. The tubes were then numbered 1-4 for ease of referencing. Four 50cm tubes were installed in each of the 6 control and eCO_2_ arrays, leading to 24 tubes in total. Data was also collected from two 2m tubes previously installed in Arrays 1 and 6. A further 50cm tube was later installed at the entrance of Array 2 for demonstrational purposes (Fig. 1D). The limited aboveground section of the tube was covered when not in use. Installation took place from 28/11/2016 to 8/12/2016, leaving 4 months for the system to equilibrate before experiments began.

### Total root imaging and physical characterisation

A Bartz Technology Corporation (VSI Bartz Technology Corporation 4187 Carpinteria Ave Unit #7. Carpinteria Ca. 93013 USA) 100X minirhizotron camera system [71] with Smucker manual indexing handle [72] was used to obtain the root images, and paired with a Bartz Technology Corporation I-CAP image capture system. The bung was carefully removed from the end of the tube, with one hand stabilising the tube during removal, and the camera was then inserted. The tubes were scanned for roots, starting from the bottom and working up. Roots are photographed and the depth, viewing angle from vertical, and tube number are recorded. The images are then analysed using SmartRoot [73] as described below, and the amount of new growth and branching is recorded through comparison with earlier images. Data was collected monthly, from all tubes in all arrays. Smart Root measurements were calibrated to cm, and, where required, volume measurements were converted to dry weight biomass using a conversion factor of 343mg dw cm^−3^, obtained from a comparison of root masses from soil cores and scans of the same samples analysed with SmartRoot (see below).

### Image analysis

Image analysis (see Figure S1) was performed using SmartRoot [73], a plugin for ImageJ [74] which we used as part of the Fiji package [75]. Roots were manually identified using the software, and a skeleton was produced with periodic width measurements along the root. The length of each root segment was recorded along with an average width across each width measurement point. Each segment was recorded with a length and width measurement in cm, along with the array number, tube number, date of sampling, depth of sampling as taken from the Smucker manual indexing handle, and angle of measurement when available. The image analysis was performed by a single person so that any subjectivity in root identification would remain constant across eCO_2_ and control populations.

### Soil coring

Soil cores were taken periodically over the two year study period; in March 2017, March, July, November of 2018, and March 2019. Three cores of length 30cm and diameter 5cm were taken from each array at BIFoR using a lined Van Walt 55mm corer. The cores were separated by horizon (visually identified by soil colour and texture) and the roots were hand-picked from each sample, and live and dead roots were separated. Live roots were identified based on the criteria in [76], with preliminary experiments to confirm this protocol using Evans Blue vital stain (Sigma Aldrich). Roots were then washed, dried, and the mass recorded for each sample.

### Root scans from soil cores

We manually recorded the length and diameter of 552 root systems (3709 roots in total) recovered from soil cores taken in March 2019. After separation and washing roots as above, roots less than 2mm thick were blotted to remove excess water and carefully teased apart using foreceps to display as much of the natural root system as possible while minimising breakages. Roots were then carefully laid out on a scanner, with as much separation as possible to aid in later image analysis (see Fig. 4A). 1cm graph paper was used as the backing to allow easy scaling of the image. The scanner lid was closed carefully to minimise movement of roots, and cleaned with a paper towel between scans to remove any debris. Each root was manually traced and automatically measured with SmartRoot.

### Birth-immigration-death model dynamics

A birth-immigration-death (BID) stochastic model can be applied to root system data by considering a unit length of root as a member of the root ‘population’. As in the main text, we take birth to represent the growth and branching of existing roots, immigration to represent a new root appearing in the minirhizotron viewing window and therefore entering the population, and death to represent the decomposition of dead roots. We will use 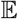 and 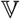 to refer to the expected value and variance of a random variable respectively. Taking *m* as the number of unit root lengths, λ*m* the birth rate, *νm* the death rate and *α* the immigration rate, this model is described by the master equation

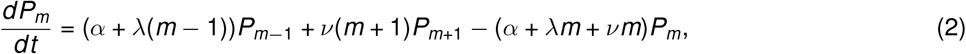

for *P_m_*(*t*), the probability of a state with *m* unit root elements at time *t*. Initial conditions at *t* = 0, 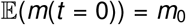 and 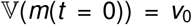, are also parameters of the model. The BID model admits a closed-form solution for an exact likelihood (which has been previously studied in stochastic biology [77]), but for simplicity and because of the continuous nature of our root observations we employ a normal approximation. Hence, we set

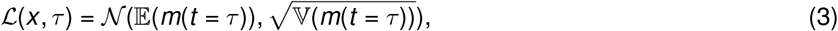

using expressions for 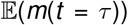 and 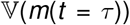, the mean and variance of root biomass observed at time *τ*, which we derive in the Supplementary Information.

### Statistical analysis and uncertainty quantification

Statistical analysis was performed in R [78] using custom scripts available at github.com/stochasticbiology/elevated-CO2. LOESS fitting was performed using the default parameterisation of the *loess* command, specifically using a span *α* of 0.75 and a polynomial degree of 2.

Caladis [50] was used for uncertainty propagation, specifically to track uncertainty through the calculation of Eqn. 5. Caladis is an online tool allowing for calculations using probability distributions [50]. Each variable in a calculation is associated with a user-defined probability distribution reflecting uncertainty in that quantity, and when a calculation is performed the value of each variable is sampled from its distribution for use in the equation. In the Supplementary Information we detail and justify the uncertainty distributions used in our NPP calculation.

### Fine root production for NPP calculation

Fine root production was calculated by observing growth of specific root branches in the top strip of a tube in each of the six arrays [24, 52, 53]. The roots were observed monthly as described above, and the growth since the month before was recorded. This allowed the calculation of total fine root production in the viewing areas for treatment and control for each of the two years of sampling. These numbers were then scaled to give a total NPP for the arrays using the geometry of the minirhizotron installations.

In the Supplementary Information we show that the minirhizotron tube samples a proportion of the total volume of the soil column in which it is embedded:

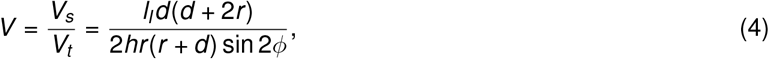

where *h* is the length of the viewing area, *r* is the radius of the minirhizotron tube, *d* is depth of viewing field, *ϕ* is the angle of the minirhizotron tube, and *l_I_* is the viewing arc length (calculate in the Supplementary Information). We further show that, given observed volume production *p_obs_* through this sampling, the total NPP estimate is given by:

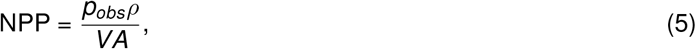

where *ρ* is root density, *A* is the area on the surface covered by the viewing area (see Supplementary Information), and *V* is calculated using equation 4.

### Raw data

All raw data (and analysis code) are available at github.com/stochasticbiology/elevated-CO2

## Competing Interests

The authors declare that no competing interests exist.

## Acknowledgements

All authors acknowledge support from the Birmingham Institute of Forest Research, and from the workshop at the School of Biosciences, University of Birmingham. IGJ acknowledges support from a Birmingham Fellowship for the University of Birmingham and Turing Fellowship from the Alan Turing Institute. BIFoR FACE is supported by the JABBS foundation and the University of Birmingham. Aleks K is supported by the Natural Environment Research Council through a CENTA studentship. This project has been supported by QUINTUS, NERC Large Grant NE/S015833/1. The authors gratefully acknowledge advice and critique received from Prof Rich Norby throughout the study.

## Author Contributions

## Supplementary Information

**Figure S1:**
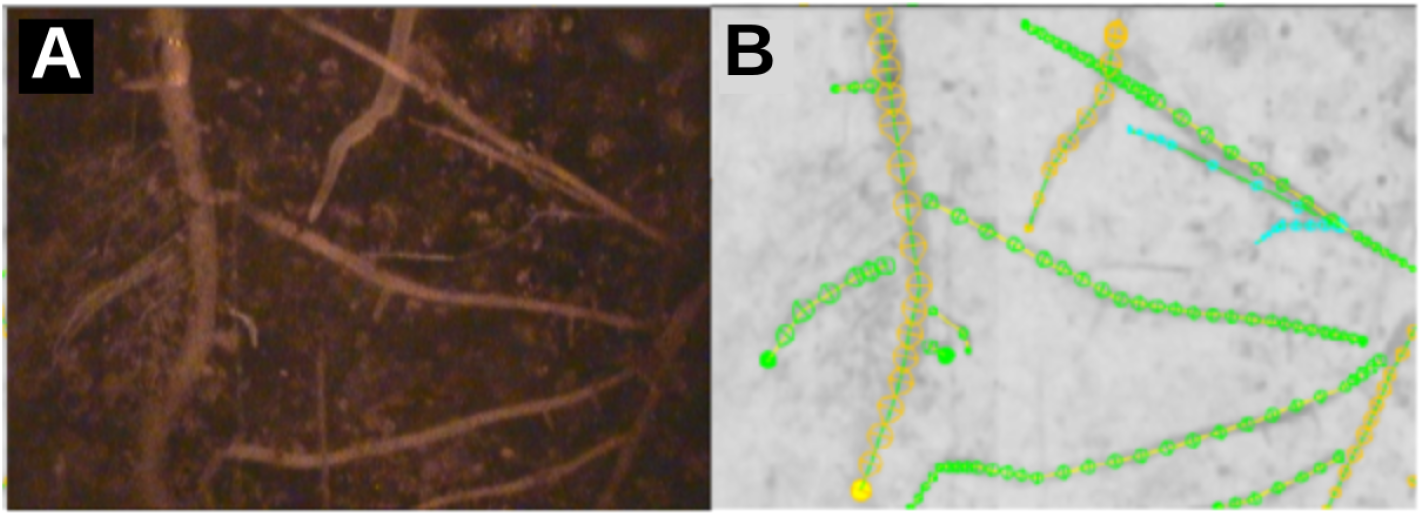
Example of image analysis perfomed on minirhizotron images. **(A)** A source image from minirhizotron sampling. **(B)** The same image following analysis with SmartRoot (see Methods). Each circle represents a root width measurement and the underlying skeleton the root length. Colours denote relative root order.

**Figure S2:**
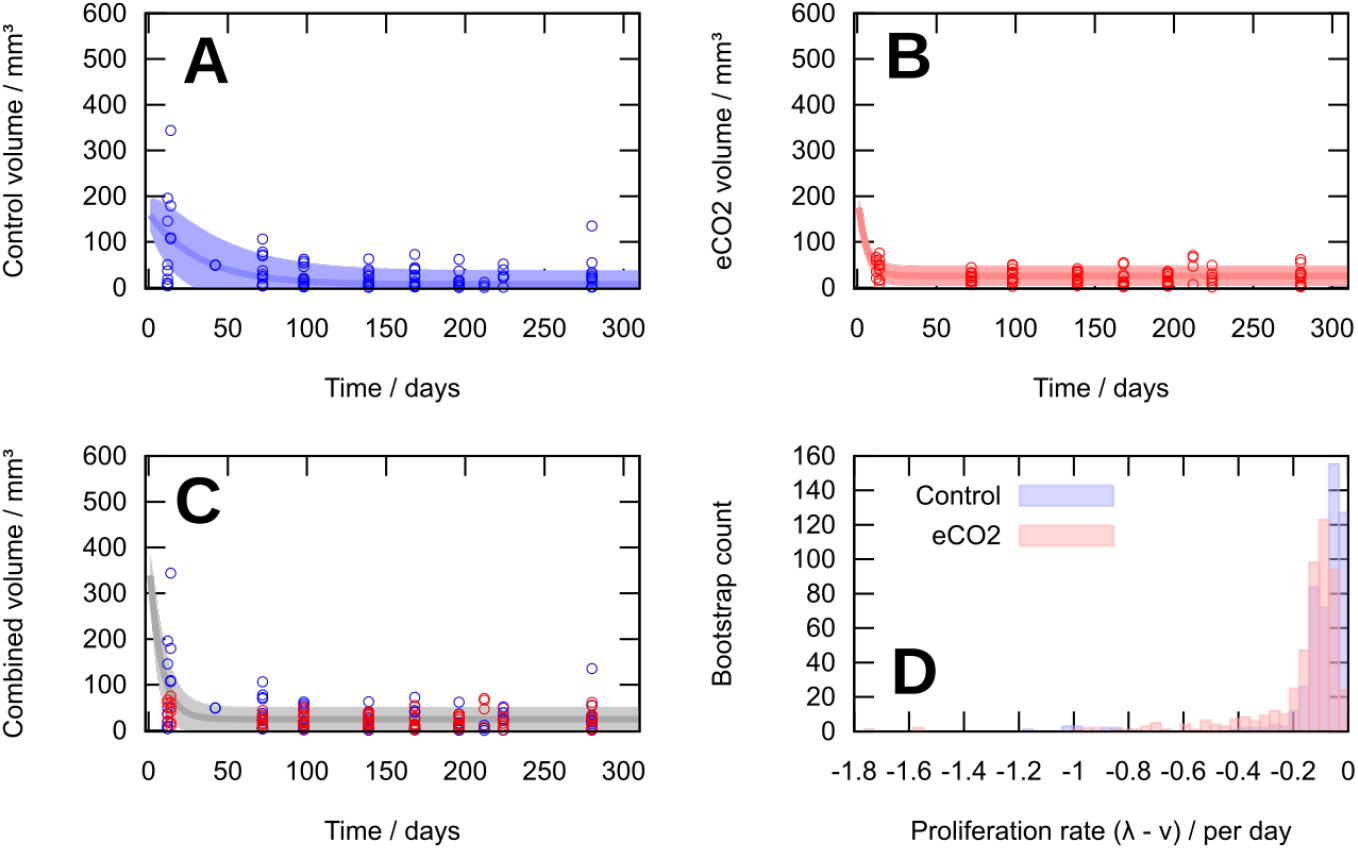
Birth-immigration-death model forsecondyearof root biomass. The birth-immigration-death (BID) model described in the text, applied to (A) eCO_2_, (B) control, and (C) combined biomass observations in year two. As in the main text, the maximum likelihood BID parameterisation is found for each dataset, then the mean and standard deviation of the model for that parameterisation is plotted. **(D)** Bootstrapped estimates for the difference between root elongation (birth, λ) and root decay (death, *ν*) parameters for eCO_2_ and control data.

### Root segment lengths from soil cores

We manually recorded the length and diameter of 552 root systems (3709 roots in total) recovered from soil cores taken in March 2019 (Fig. 3D). We found an increase in root segment diameter length in eCO2 versus control (Fig. S3) (see Results). However, the length result should be interpreted with some caution, as the root segment length recorded from a soil core scan may not be representative of the root length belowground. Roots are often cut in the soil coring process, and many fine roots break during removal from the soil, in the washing process, and during separation for scanning (see Methods). This has lesser impact on biomass and width measurements as long as these segments are preserved, however this can affect the recorded lengths – however, there is no reason that this effect would systematically affect control and treatment experiments diffeently, so identified differences are likely still representative.

**Figure S3:**
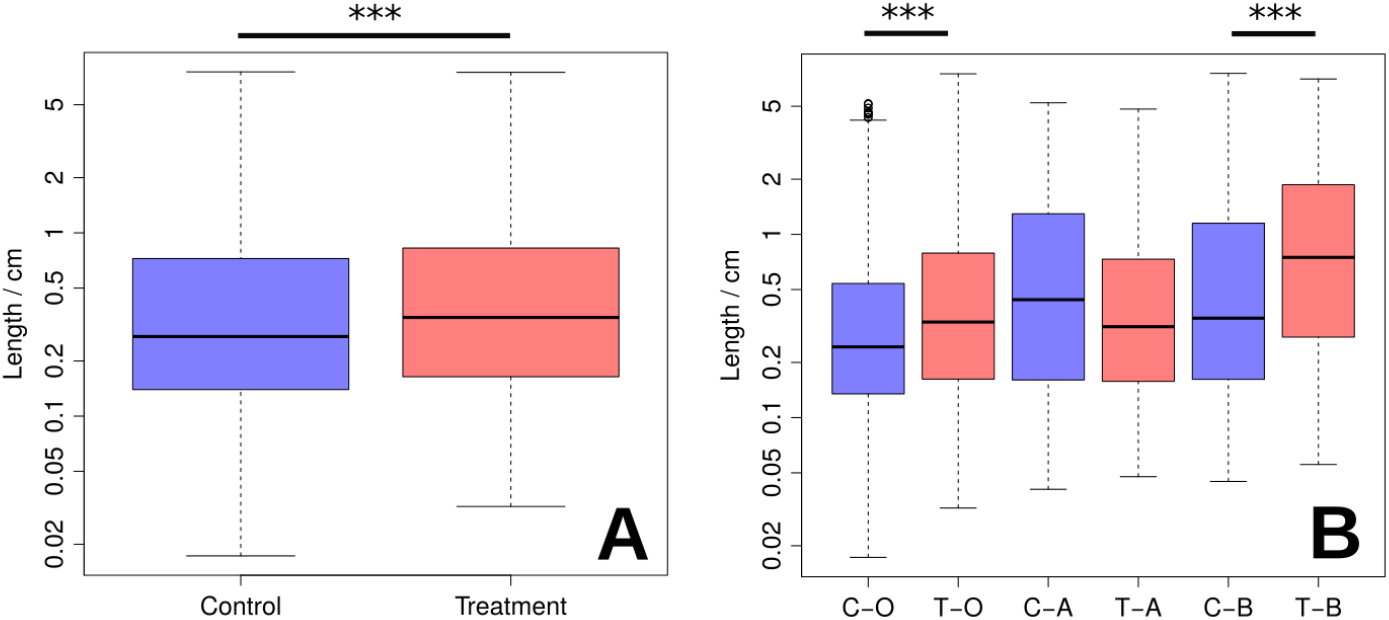
Root lengths sampled from soil cores. Length of live root segments taken from March 2019 soil cores under eCO_2_ (red) and control (blue) conditions. **(A)** Logged length of live roots separated by treatment and control. A Mann Whitney test shows a significant increase in root segment length under eCO_2_ (1.11-fold, *p* = 1.7 × 10^−6^). **(B)** Logged root lengths by horizon for eCO_2_ (red) and control (blue). The increase in length under eCO_2_ is consistent across the O and B horizons but not visible in the A horizon (1.21-fold, *p* = 9.9 × 10^−10^; 0.76-fold, *p* = 0.078; 1.52-fold, *p* = 8.9 × 10^−4^ for O, A, and B respectively).

### BID statistics

We will use 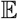 and 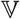 to refer to the expected value and variance of a random variable respectively. We start with the BID master equation, describing the probability *P*(*m, t*) of observing *m* length elements at time *t* under the influence of immigration *α*, birth λ and death *ν*:

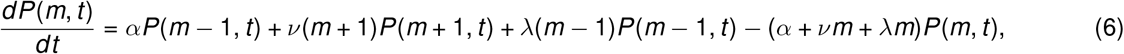

with initial condition enforcing that *m* = *m*_0_ at *t* = 0:

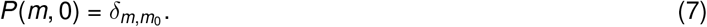

Defining the generating function 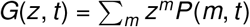, we obtain the following PDE from Eqn. 6

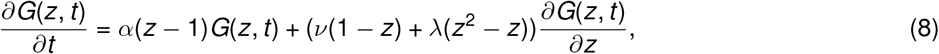

with initial condition

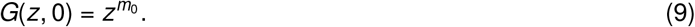

The solution is readily found through the method of characteristics [77]:

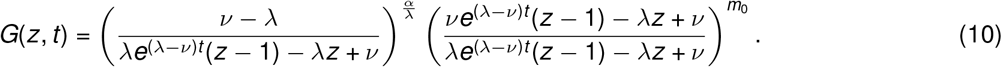

We can obtain necessary information about *P*(*m, t*) from the generating function (Eqn. 10). *P*(*m*) is given by

**Figure S4:**
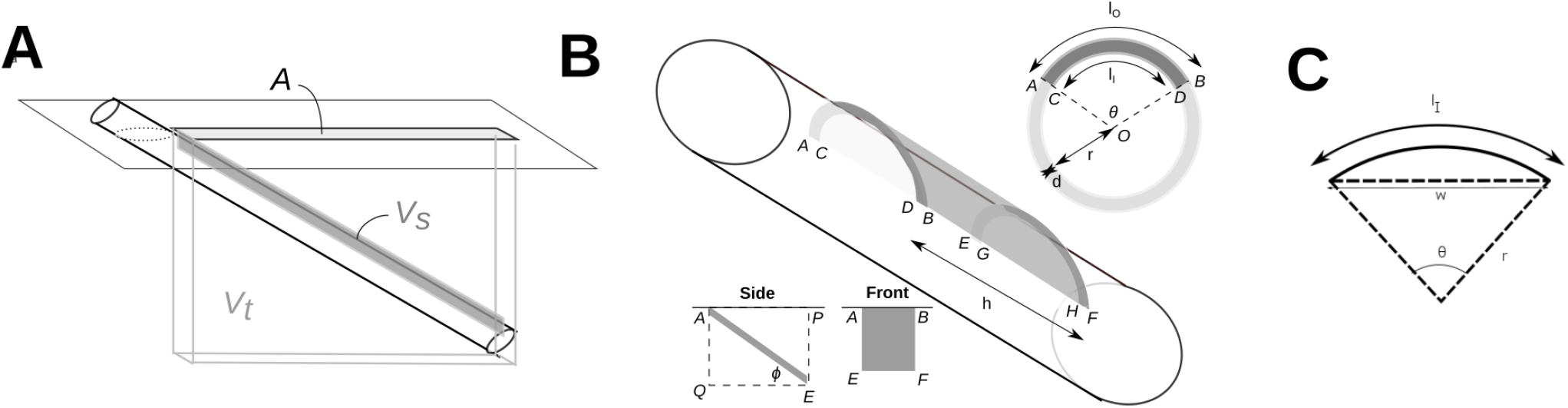
Model geometry for production calculations. **(A)** A minirhizotron tube in soil, with sampling volume *V_s_* a subset of total volume *V_t_*, which is projected to the soil surface *A*. **(B)** Geometry and dimensions for derivations in the text. **(C)** The image width, *w*, in relation to the viewing arc, *l_I_*, the viewing angle, *θ*, and the radius, *r*.

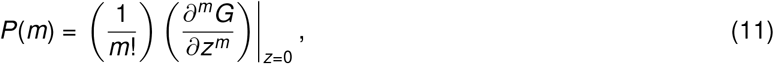

the expected value by

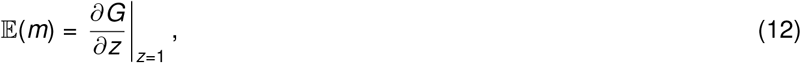

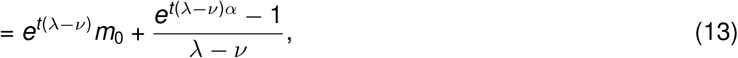

and the variance by

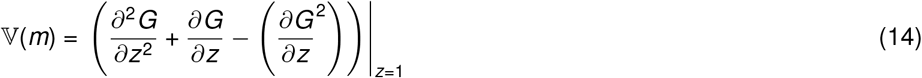

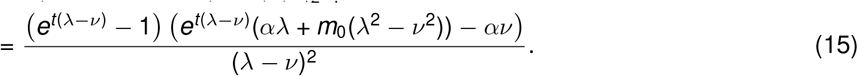

Note that for the purposes of this model we add an additional value *v*_0_ to the variance. This linear noise contribution accounts for experimental variance due to noisy observations:

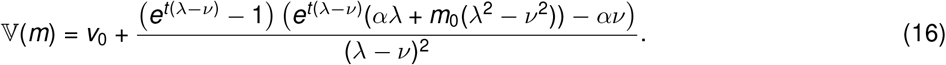

### NPP calculation

#### Calculating fine root NPP from root production values

We model a rhizotron tube *in situ* at an angle *ϕ* with the horizontal soil surface. Root images are collected covering a segment with angle *θ*, and corresponding image width *w*, which allows a viewing window of width *l_I_* with depth of field, *d*, as shown in Figure S4. We aim to produce a 2D projection, *A*, of the volume sampled by the minirhizotron, *V_s_*, onto the surface for the estimation of root NPP as a production per unit area. To do this, we define the proportion of the total volume below this region (*A*) that has been sampled using the minirhizotron, and call this volume *V_t_* (Figure S4A).

The viewing angle, *θ* (in radians) can be calculated from the viewing arc *l_I_* and the tube radius *r*

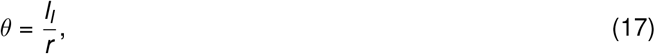

and therefore the outer arc *l_O_* can be determined from the depth of field (Figure S4B):

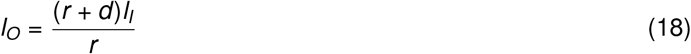

The sampling volume around the minirhizotron, *V_s_*, is *ABCDEFGH* in Figure S4B. From the area of a circle sector, area *ABO* is 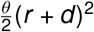 and area *CDO* is 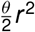, so area *ABCD* is 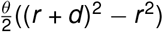 and

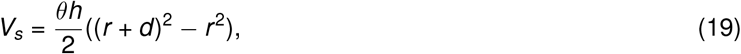

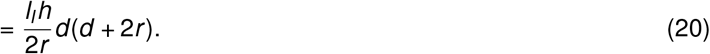

The total volume *V_t_* is delimited by the extreme points of the sampling volume *ABEFPQ*. From Figure S4B it is readily seen that lengths *AE* = *h*cos *ϕ*, *AQ* = *h*sin *ϕ*, *AB* = (*r* + *d*), so

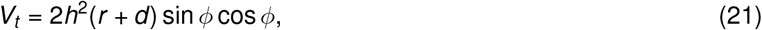

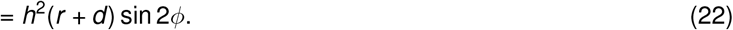

Using equations 20 and 22, we obtain the proportion of the total volume sampled by the minirhizotron:

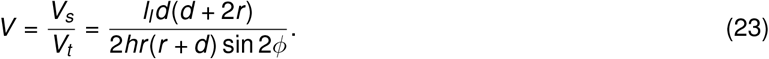

#### Obtaining the sampled arc length *l_I_* from the image width

The viewing arc, *l_I_*, can be calculated from the image width, *w*, and the tube radius, *r* (Figure S4C), using the following set of equations:

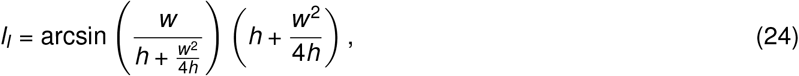

where

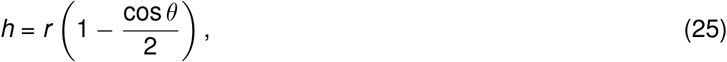

and

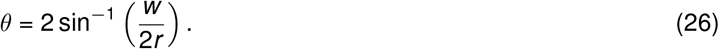

#### Scaling to total fine root NPP

Let *p* be the observed production, calculated as *p* = *p_obs_ρ*, where *p_obs_* is observed increase in root volume and *ρ* is the density of roots. We can estimate total NPP as:

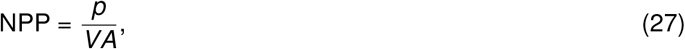

where *A* is the area at the surface that has been imaged (see Figure S4).

In this case *A* is given by

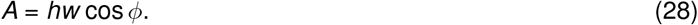

where *h* is the length and *w* the width of the viewing area and *ϕ* is the angle of the tube (see Figure S4).

### Calculating NPP using Caladis

Caladis allows for calculations using probability distributions [50]. Each variable is associated with a user-defined probability distribution, and when a calculation is performed the value of each variable is first pulled from its distribution for use in the equation. By repeating the calculation with a new set of variables each time, a histogram is produced showing the result of the calculation after a default of 20000 iterations.

Our NPP calculation uses the following equation (derived above):

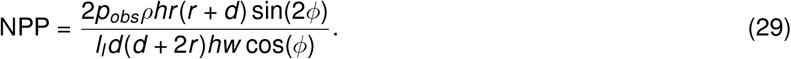

Each variable in equation 29 has an associated probability distribution, defined below. Unless otherwise stated, *σ_x_* is chosen to reflect a 95% confidence level within 2*σ_x_* of the mean.

- 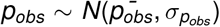, where 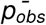 is the total observed root volume increase (cm^−3^) and *σ_p_obs__* is the standard deviation of this increase for each array, calculated from our data (values below).
- 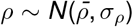, where 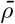 is mean root density (g dw cm^−3^) and *σ_ρ_* is the standard deviation of this density. Our calculation used 0.343 ± 0.161 g dw cm^−3^ calculated from our data.
- 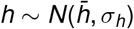, where 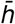 is the total viewing length along the rhizotron (m) (see Methods). Our calculation used *h* ~ *N*(0.1755,0.005) to allow for inacurracies in the measurement of the viewing windows.
- 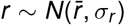, where 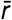 is the the radius of the minirhizotron tube (m). Our calculation used *r* ~ *N*(0.0275,0.005) to allow for variations in tube manufacturing.
- *d* ~ *U*(*d*_−_, *d*_+_), where *d*_−_ and *d*_+_ represent the maximum and minimum depth of field values (m). These values were chosen to represent the spread of depth of field values in [68], centred around the standard value used. A uniform distribution *d* ~ *U*(0.0005,0.0035) was used to reflect the uncertainty as explained in [68], rather than this being a measurement.
- 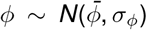, where 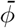 is the angle of the minirhizotron tube. Our calculation used *ϕ* ~ *N*(0.785,0.196) (radians) to cover variations in tube installation.
- 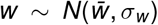, where 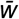 is the width of the viewing area. Our calculation used *l_i_* ~ *N*(0.0189,0.005) to capture inaccuracies in the measurement and calculation of this width.
- 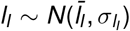, where 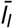 is the viewing arc length (m) (see Methods). Our calculation used *l_i_* ~ *N*(0.0192,0.005) to capture inaccuracies in the measurement and calculation of this arc length.

Links to the Caladis calculations for each case are:

- Control Y1 (*p_obs_* ~ *N*(0.0246,0.0196)) http://www.caladis.org/compute/?q=2*%24pobs*%24rho*%24h*%24r*(%24r%2B%24d)*sin(2*%24phi)%2F(%24l*%24d*(%24d%2B2*%24r)*%24h*%24w*cos(%24phi))&v=pobs%3Anorm%2C0.0246%2C0.0196%3Brho%3Anorm%2C0.343%2C0.161%3Bh%3Anorm%2C0.1755%2C0.005%3Br%3Anorm%2C0.0275%2C0.005%3Br%3Anorm%2C0.0275%2C0.005%3Bd%3Aunif%2C0.0005%2C0.0035%3Bphi%3Anorm%2C0.785%2C0.196%3Bl%3Anorm%2C0.0192%2C0.005%3Bd%3Aunif%2C0.0005%2C0.0035%3Bd%3Aunif%2C0.0005%2C0.0035%3Br%3Anorm%2C0.0275%2C0.005%3Bh%3Anorm%2C0.1755%2C0.005%3Bw%3Anorm%2C0.0189%2C0.005%3Bphi%3Anorm%2C0.785%2C0.196&x=off&n=m&h=fd&a=rad
- Control Y2 (*p_obs_* ~ *N*(0.00737,0.00319)) http://www.caladis.org/compute/?q=2*%24pobs*%24rho*%24h*%24r*(%24r%2B%24d)*sin(2*%24phi)%2F(%24l*%24d*(%24d%2B2*%24r)*%24h*%24w*cos(%24phi))&v=pobs%3Anorm%2C0.00737%2C0.00319%3Brho%3Anorm%2C0.343%2C0.161%3Bh%3Anorm%2C0.1755%2C0.005%3Br%3Anorm%2C0.0275%2C0.005%3Br%3Anorm%2C0.0275%2C0.005%3Bd%3Aunif%2C0.0005%2C0.0035%3Bphi%3Anorm%2C0.785%2C0.196%3Bl%3Anorm%2C0.0192%2C0.005%3Bd%3Aunif%2C0.0005%2C0.0035%3Bd%3Aunif%2C0.0005%2C0.0035%3Br%3Anorm%2C0.0275%2C0.005%3Bh%3Anorm%2C0.1755%2C0.005%3Bw%3Anorm%2C0.0189%2C0.005%3Bphi%3Anorm%2C0.785%2C0.196&x=off&n=m&h=fd&a=rad
- Treatment Y1 (*p_obs_* ~ *N*(0.0290,0.0153)) http://www.caladis.org/compute/?q=2*%24pobs*%24rho*%24h*%24r*(%24r%2B%24d)*sin(2*%24phi)%2F(%24l*%24d*(%24d%2B2*%24r)*%24h*%24w*cos(%24phi))&v=pobs%3Anorm%2C0.0290%2C0.0153%3Brho%3Anorm%2C0.343%2C0.161%3Bh%3Anorm%2C0.1755%2C0.005%3Br%3Anorm%2C0.0275%2C0.005%3Br%3Anorm%2C0.0275%2C0.005%3Bd%3Aunif%2C0.0005%2C0.0035%3Bphi%3Anorm%2C0.785%2C0.196%3Bl%3Anorm%2C0.0192%2C0.005%3Bd%3Aunif%2C0.0005%2C0.0035%3Bd%3Aunif%2C0.0005%2C0.0035%3Br%3Anorm%2C0.0275%2C0.005%3Bh%3Anorm%2C0.1755%2C0.005%3Bw%3Anorm%2C0.0189%2C0.005%3Bphi%3Anorm%2C0.785%2C0.196&x=off&n=m&h=fd&a=rad
- Treatment Y2 (*p_obs_* ~ *N*(0.0108,0.00485)) http://www.caladis.org/compute/?q=2*%24pobs*%24rho*%24h*%24r*(%24r%2B%24d)*sin(2*%24phi)%2F(%24l*%24d*(%24d%2B2*%24r)*%24h*%24w*cos(%24phi))&v=pobs%3Anorm%2C0.0108%2C0.00485%3Brho%3Anorm%2C0.343%2C0.161%3Bh%3Anorm%2C0.1755%2C0.005%3Br%3Anorm%2C0.0275%2C0.005%3Br%3Anorm%2C0.0275%2C0.005%3Bd%3Aunif%2C0.0005%2C0.0035%3Bphi%3Anorm%2C0.785%2C0.196%3Bl%3Anorm%2C0.0192%2C0.005%3Bd%3Aunif%2C0.0005%2C0.0035%3Bd%3Aunif%2C0.0005%2C0.0035%3Br%3Anorm%2C0.0275%2C0.005%3Bh%3Anorm%2C0.1755%2C0.005%3Bw%3Anorm%2C0.0189%2C0.005%3Bphi%3Anorm%2C0.785%2C0.196&x=off&n=m&h=fd&a=rad

### Raw data

Minirhizotron measurements, soil core measurements, root morphological measurements, raw production observations, and Caladis outputs are available as raw data files from github.com/stochasticbiology/elevated-CO2

## Notes

https://github.com/StochasticBiology/elevated-co2/

### Competing Interest Statement

The authors have declared no competing interest.

